## Supplemental Figure 1, 2 and 3 for "Astrocytes mediate two forms of spike timing-dependent depression at entorhinal cortex-hippocampal synapses"

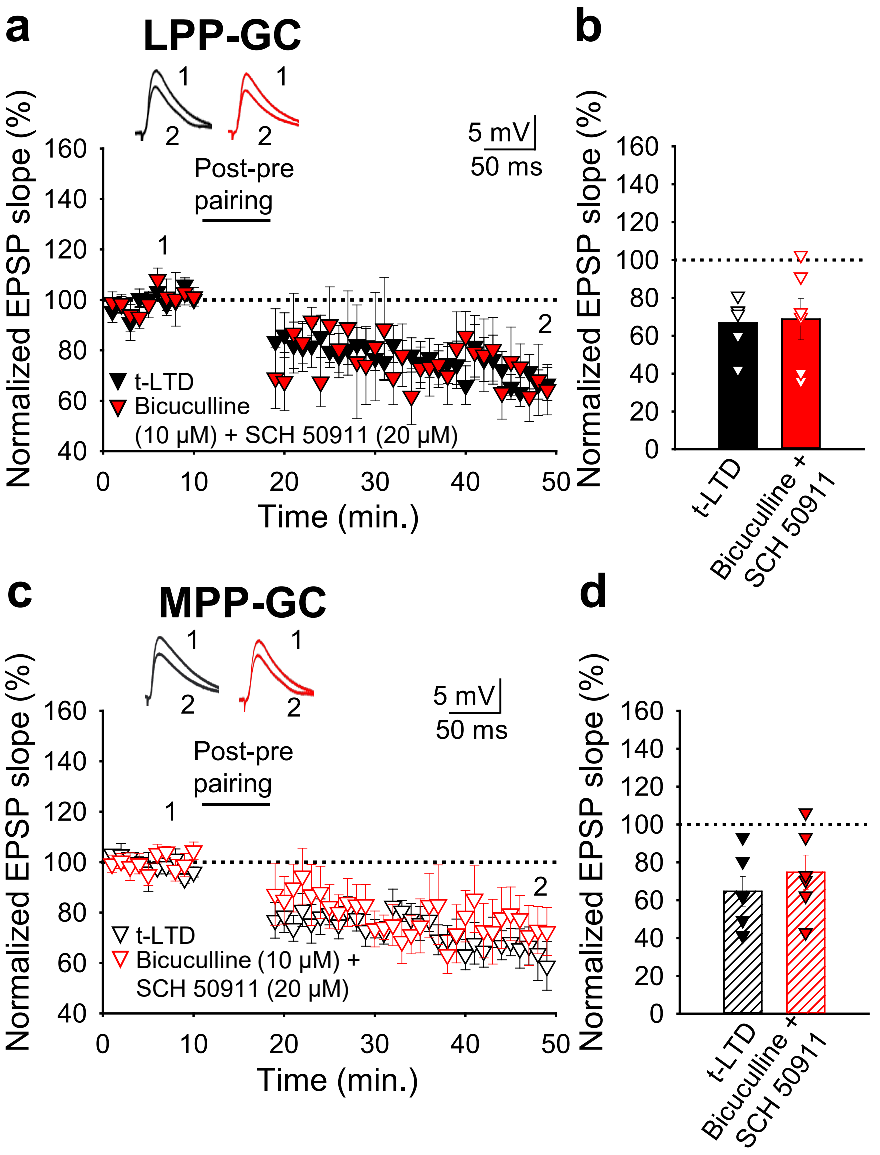


**Supplementary Figure 1. GABA receptors are not required for t-LTD at LPP- or MPP-GC synapses.** (A) and (C) Time course of EPSP monitored in slices during baseline and after the addition of bicuculline (LPP-GC: 67 ± 6%, n = 6; MPP-GC: 65 ± 8%, n = 6) and SCH50911 (LPP-GC: 69 ± 11, n = 6; MPP-GC: 75 ± 9%, n = 6) to the perfusion fluid. Traces show the EPSP before (1) and 30 min after (2) treatment. (B) Summary of the results. Error bars indicate S.E.M.

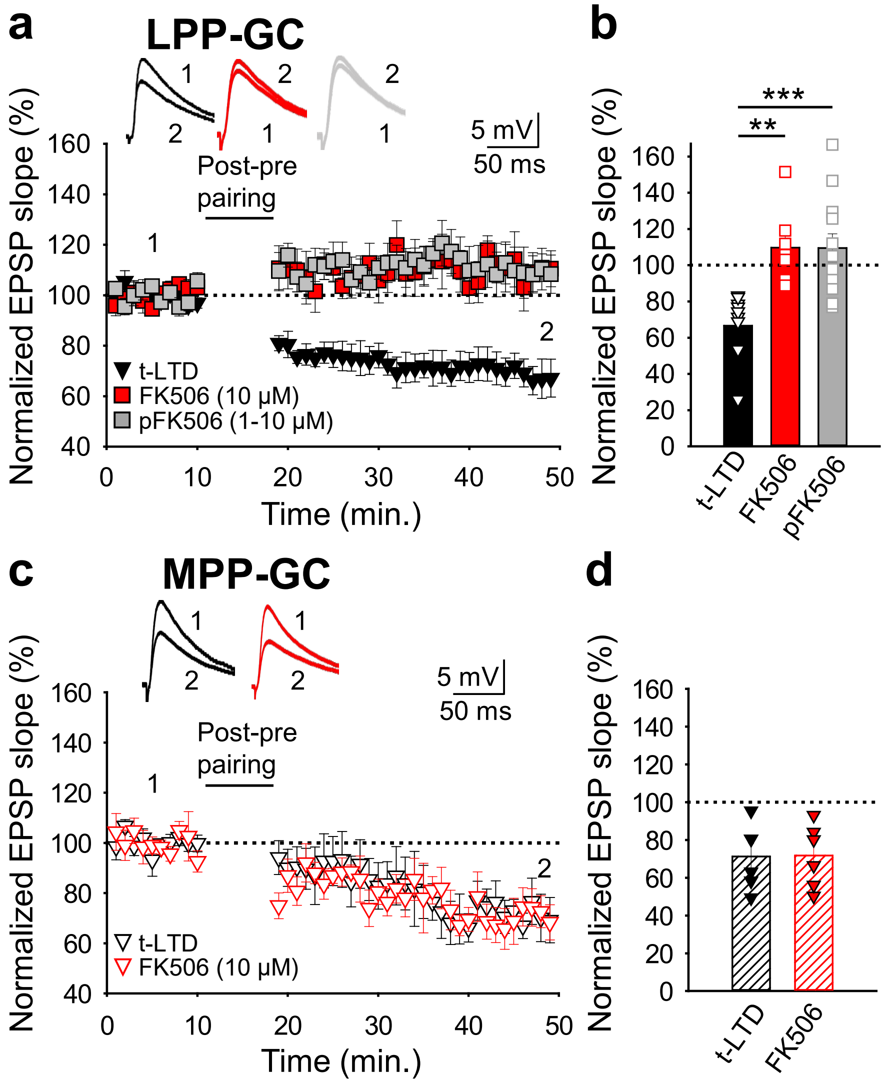


**Supplementary Figure 2. Calcineurin is required for t-LTD induction at the LPP, but not at MPP-GC synapses.** (**a**) t-LTD requires postsynaptic calcineurin at LPP-GC synapses. The EPSP slopes monitored in control slices (black triangles, n = 9) and in slices treated with the calcineurin blocker FK506 (1-10 μM), either in the bath (red squares, n = 8) or loaded into the postsynaptic neuron (gray squares, n = 13) are shown. Traces show EPSP before (1) and 30 min after (2) pairing. (**b**) Summary of the results. (**c**) t-LTD does not require calcineurin at MPP-GC synapses. The EPSP slopes monitored in control slices (open black triangles, n = 6) and in slices treated with FK506 (10 μM) in the bath (open-red triangles, n = 6) are shown. Traces show EPSP before (1) and 30 min after (2) pairing. (**d**) Summary of the results. $**p < 0.01$, One-way ANOVA + Holm–Sidak. Error bars represent the S.E.M.

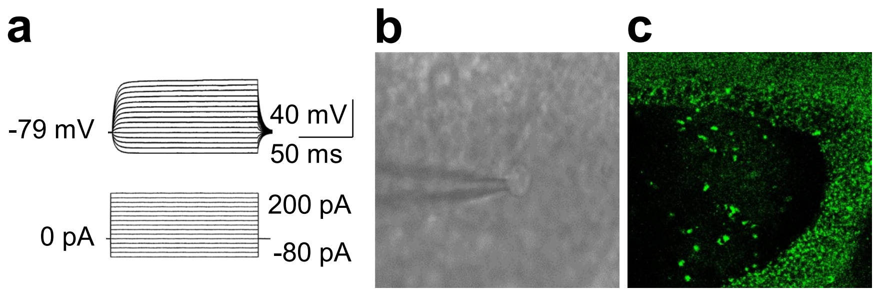


**Supplementary Figure 3. Identification of astrocytes**. **(a)** Low magnification infrared video microscopy image showing a brain slice containing LPP- and MPP-synapses. Stimulating (S) and recording (R) electrodes are indicated as well as an astrocyte. **(b)** Voltage responses of typical astrocytes in the surrounding of LPP-and MPP-GC synapses are shown. Astrocytes show a passive response to current injections. **(c)** 25x confocal image showing the GFP fluorescence in the granule cell layer of the dentate gyrus from a dnSNARE mouse off Dox since birth. No GFP could be found in neurons.
